# FSTL1 is an antagonist of ERK1/2 phosphorylation during ciliogenesis and preadipocyte differentiation

**DOI:** 10.1101/2024.02.21.581425

**Authors:** Leonardo Santos, Lucia Guggeri, Carlos Escande, José Luis Badano, Victoria Prieto-Echagüe

## Abstract

FSTL1 is a secreted glycoprotein that is involved in several processes in health and disease, including development, cardiovascular disease, cancer, inflammation, and obesity. The signaling pathways used by FSTL1 to act on target tissues seem to activate different intracellular mediators specific to each tissue and several of the mechanisms of action remain to be determined at the molecular level, including intracellular mediators and receptors. We have previously unveiled a novel role for FSTL1 in ciliogenesis and provided evidence for an Fstl1/cilia axis in preadipocyte differentiation. This pathway is relevant to the pathogenesis of obesity and of a group of conditions called ciliopathies since they are caused by the dysfunction of the primary cilia. This work aimed to identify intracellular mediators of FSTL1 action on ciliogenesis and adipogenesis. We analyzed ERK phosphorylation levels as well as cilia length in the absence of FSTL1 and in the presence of the pERK inhibitor U0126. We also analyzed the differentiation and cilia dynamics of U0126-treated preadipocytes and tested the ERK-mediated signaling by BMP4 in the presence of added extracellular Fstl1. Here, we propose that MAP kinase ERK is a mediator of ciliogenesis downstream of FSTL1 and provide additional data that suggest that FSTL1 antagonizes BMP non-canonical signaling to modulate ciliogenesis and adipogenesis. In sum, our data reinforce the interest on the axis FSTL1/cilia in the modulation of adipogenesis and provide evidence to add ERK to this working model.

## Introduction

FSTL1 is a secreted glycoprotein that plays a role in several processes in health and disease (reviewed in [1]). FSTL1 participates in normal developmental processes such as in lung maturation, since Fstl1 KO mice die at birth due to respiratory malfunction caused by defective lung organogenesis [2, 3]. FSTL1 has also been linked to the pathogenesis of several conditions including cardiovascular disease showing a protective role in myocardial infarction [4, 5]. In addition, FSTL1 has been studied in cancer although its role in this process remains unclear due to contradictory evidence found in different experimental models. As non-exhaustive examples, in ovarian and endometrial cancer, FSTL1 was proposed to be a tumor suppressor [6, 7], while in glioblastoma FSTL1 is overexpressed compared to normal tissue [8] and a metastatic brain cancer cell line has a higher FSTL1 expression compared to its primary breast cancer cell line [9]. The role of FSTL1 in inflammation and immune disease is also controversial with reports of both anti- and pro-inflammatory effects [1]. For instance, FSTL1 was shown to have a pro-inflammatory role in a septic shock mouse model where it promotes the secretion of IL1beta by monocytes/macrophages [10]. Conversely, FSTL1 downregulates IL-1beta in a cisplatin nephrotoxicity model [11].

FSTL1 also plays a role in adipogenesis and obesity. It was first proposed as a biomarker for adipogenesis given that it is expressed in pre-adipocytes, but its expression is significantly reduced in mature adipocytes [12]. More recently we were able to demonstrate that FSTL1 plays an active role in the differentiation of 3T3-L1 preadipocytes. We showed that both high levels of FSTL1 at the onset and its subsequent reduction during the process are required for normal differentiation in this cell line [13]. Importantly, in vivo experiments by others confirmed that differentiation to adipocytes of Fstl1-/- mouse embryonic fibroblasts (MEFs) and stromal vascular cells (SVCs) was impaired [14]. Moreover, different reports are starting to link FSTL1 with obesity in humans (see for example [15, 16]).

Despite the abundance of evidence implicating FSTL1 in diverse biological processes and signaling pathways, the mechanism of action of FSTL1 remains to be determined at the molecular level, including the receptors and intracellular mediators involved in each model. FSTL1 was first cloned as a member of the TGFβ family as an TGFβ1 inducible gene [17, 18]. Later, FSTL1 was shown to bind to members of this family, namely Activin A, ActR-IIB, BMP4, BMPR1A, TGFβ1 [19, 20]. Also, in a model of overexpression in mammalian cells FSTL1 was shown to bind to BMP4 and to interfere with BMP4/BMPRII downstream signaling pathway mediated by SMAD1/5/9 [3]. Structurally, FSTL1 comprises a N-terminal signal secretion peptide; a Follistatin-like domain, a Kazal-like domain, two EF-hand domains and a VWFC domain (UniProt:Q12841) and this domain organization might influence its binding partners or target molecules [21].

The signaling pathways used by FSTL1 to exert its action on target tissues have been studied in different models and these pathways seem to differ in different organs, binding to different receptors and activating different intracellular mediators specific to each tissue or model. For instance, in cardiac myocytes, the protein DIP2a was proposed as a receptor of FSTL1 in a signaling pathway mediated by the phosphorylation of AKT in the context of the cardiovascular protective role of FSTL1 [22]. Although binding partners such as DIP2A, TLR4, BMP receptors and intracellular mediators have been shown to act downstream of extracellular FSTL1, others might probably exist [1].

In our previous work describing the role of FSTL1 in the differentiation of 3T3-L1 we showed that FSTL1 is a novel regulator of primary cilia: knocking down FSTL1 in hTert-RPE1 cells decreases primary cilia length while adding Fstl1 back in the cell culture media rescues the ciliary phenotype [13]. Cilia are antennae-like organelles that are present in virtually every cell in vertebrates and act as signaling hubs [23]. Furthermore, defects in ciliary structure or function have been described to be causally associated to a set of human pathologies called ciliopathies [23]. Cilia could be regarded as a downstream target of FSTL1 and thus understanding how FSTL1 regulates cilia is critical to dissect its function both in homeostatic and pathological conditions. For example, the primary cilium and ciliary proteins have been shown to be regulators of adipogenesis (reviewed in [24]). During adipocyte differentiation cilia show a similar pattern compared to FSTL1: cilia are present in pre-adipocytes but are reabsorbed as differentiation proceeds [13, 25]. We showed that downregulating FSTL1 in 3T3-L1 cells resulted in shortened cilia at the onset of differentiation which in turn inhibited differentiation. Importantly, artificially maintaining FSTL1 levels up along differentiation impaired the naturally occurring cilia shortening and resulted in differentiation inhibition [13]. FSTL1 could therefore regulate pathways relevant to adipogenesis directly and/or through the cilium. Some of the signaling pathways coordinated by primary cilia are activated or mediated by Ca2+, Notch, receptor Tyrosine kinase receptors such as PDGF, EGFR, IGF1R. TGFβ-BMP has also been associated with the primary cilium and cilia modulates Wnt, and Hedgehog, two anti-adipogenic signaling cascades (reviewed in [26].

The aim of this work was to identify the signaling pathway that mediates the modulation of cilia length by FSTL1 in hTert-RPE1 cells, including intracellular mediators and possibly a plasma membrane receptor. We hypothesized that proteins belonging to the TGFβ family are involved as receptors or intracellular mediators and investigated both canonical and non-canonical signaling in response to FSTL1 knock-down. We show here that FSTL1 regulates ERK activation, and that cilia length regulation is mediated by ERK phosphorylation in human hTERT-RPE1 and murine 3T3-L1 cells. Moreover, we show that antagonizing BMP4 signaling by FSTL1 inhibits the non-canonical pathway mediated by ERK phosphorylation. Taken together, our data suggest that FSTL1 regulates cilia length by potentially binding to a receptor of the BMP family thereby modulating BMP4 signaling, implicating potential roles for BMP4 and its receptors in the pathogenesis of ciliopathies.

## Results

### 1. FSTL1 modulates phospho-ERK in hTERT-RPE1 cells

As mentioned, the goal of this work was to gain insight into the mechanism by which FSTL1 regulates cilia length. Based on the literature, one strong candidate was the MAP kinases cascade and ERK in particular since this central pathway has been shown to regulate ciliogenesis [27]. For example, dephosphorylation of ERK by DUSP6 phosphatase reduces cilia length in RPE cells and in Chlamydomonas reinhardtii [27]. Also, cisplatin treated cells kidney tubular epithelial cells have shorter cilia and activated ERK while inhibition of pERK by U0126 during cisplatin treatment elongates cilia [28]. Importantly, FSTL1 was shown to bind to members of TGFβ family BMP4 and to antagonize pSMAD signaling by BMP4 [3]. In turn, BMP4 canonical signaling cross talks with other pathways including MAPK pathway [29, 30]. Moreover, ERK phosphorylation has been previously shown to participate in FSTL1 signaling in ways that might be tissue-dependent. For instance, in cardiomyocytes and cardiac fibroblasts, cell survival to hypoxia/reperfusion stress is favored by Fstl1-promoted upregulation of pErk [5, 31]. On the other hand, in a Fstl1^+/-^ mouse model of pulmonary hypertension (HPH) addition of FTSL1 attenuates the symptoms of the disease by downregulating pERK in pulmonary artery smooth muscle cells and thereby preventing fibrosis [32].

To test whether ERK mediates the effects of FSTL1 on ciliogenesis we reduced the expression of FSTL1 using our previously validated siRNA [13] and analyzed the phosphorylation of the MAP Kinase ERK. Downregulation of FSTL1 in the same conditions that induce ciliogenesis (cell culture confluence and 24h-serum starvation) induces an average of a 2-fold upregulation of pERK as a significant increase detected by Western blot (Fig. 1A).

**Figure 1.**
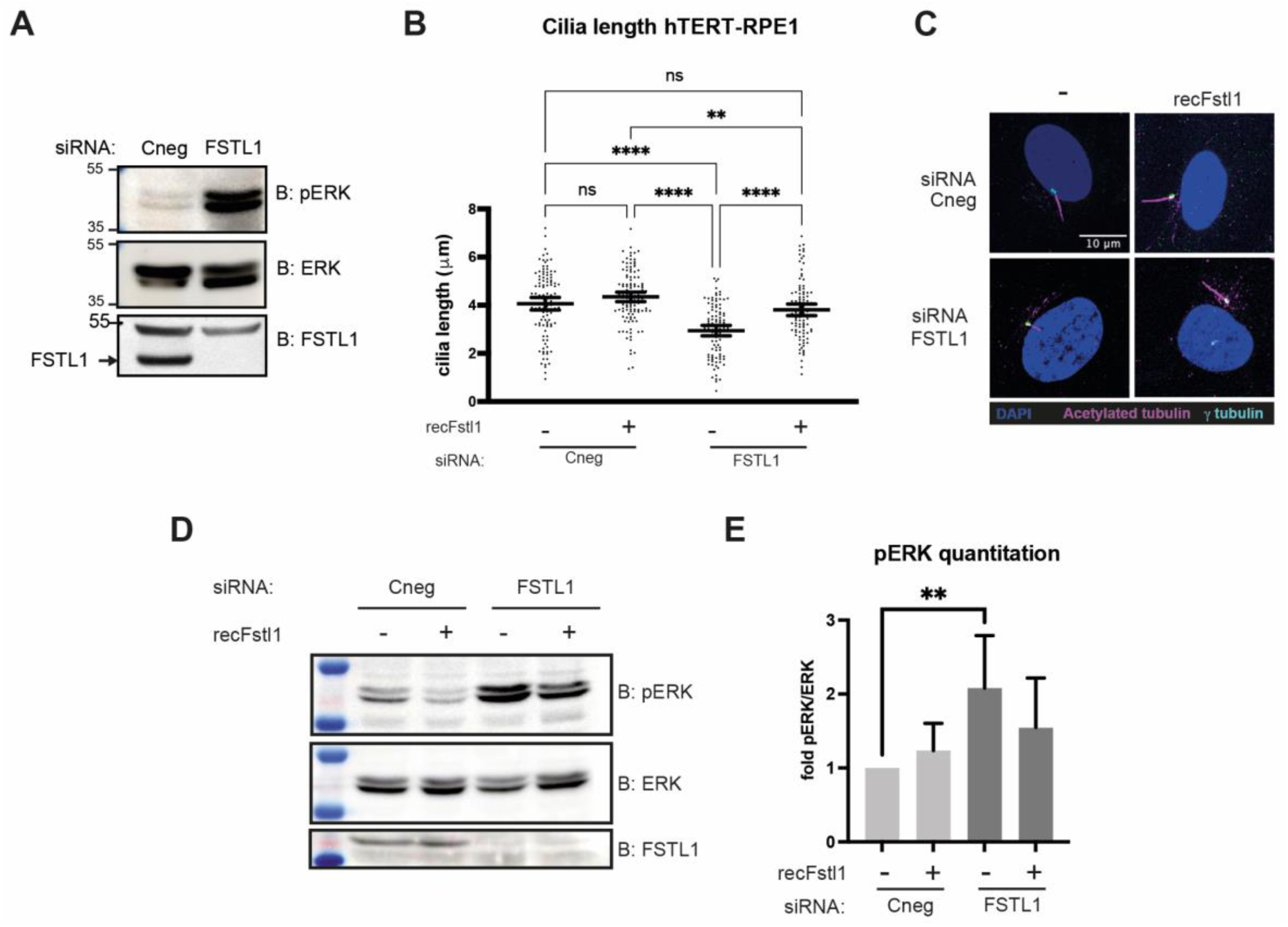
FSTL1 KD activates ERK phosphorylation in hTERT-RPE1 cells and extracellular Fstl1 reverses ERK phosphorylation and cilia length. (A) hTERT-RPE1 cells were transfected with siRNA Cneg or siRNA FSTL1 for 48 hours. In the last 24 hours FBS was removed and cell lysates were analyzed by Western blot anti-pERK, anti-ERK and anti-FSTL1. (B-E) hTERT-RPE1 cells were transfected with siRNA Cneg or siRNA FSTL1 for 48 hours. In the last 24 hours cell culture media were changed and FBS-medium was added with or without supplementation with 200 ng/ml recFstl1. (B) coverslips from the same experiment shown in (D) were immunostained for acetylated tubulin and gamma-tubulin and primary cilia were measured in each condition. (C) micrographs showing basal bodies (γ-tubulin) in *cyan*, cilia (acetylated tubulin) in *red* and nuclei in *blue* (DAPI). Scale bar represents 10 μm. Cilia lengths from each condition were compared using a Kruskall-Wallis test. Dot plots with a line at the mean are shown and error bars represent 95% confidence intervals. (D) cell lysates were analyzed by Western blot anti-pERK, anti-ERK and anti-FSTL1. (E) Densitometry readings were used to calculate pERK/ERK ratios with each condition normalized to the siRNA Cneg transfected and untreated cells. Data pooled from 10 experiments are shown in the bar graph and error bars represent 95% confidence intervals. Significant differences are signaled with **: P=0.001-0.01; ***: P=0.0001-0.001; ****: P<0.0001. Data in A and C are representative of 3 and 2 experiments respectively.

We had previously shown that at least part of the effect of FSTL1 on cilia regulation is cell-non-autonomous relying on secreted FSTL1 acting in a paracrine manner since adding recombinant Fstl1 to the extracellular media was able to rescue the cilia phenotype of FSTL1 knockdown cells [13]. We first recapitulated these results (Fig. 1B-C) and then we analyzed ERK phosphorylation levels in each condition. FSTL1 KD decreases cilia length from 4.06 μm to 2.95 μm (Fig. 1B) and increases ERK phosphorylation (Fig. 1A and 1D) compared to control cells transfected with a scramble siRNA. In turn, adding recombinant Fstl1 to the extracellular media of FSTL1-KD cells was able to rescue the cilia length to 3.81 μm and to decrease ERK phosphorylation compared to control cells and to untreated FSTL1-KD cells. Thus, incubation of FSTL1 depleted cells with recombinant extracellular FSTL1 rescues not only the shorter cilia phenotype but also the phosphorylation of ERK. Therefore, our data show a correlation between shorter cilia length and higher ERK phosphorylation and suggest that the regulation of cilia length by FSTL1 is mediated by ERK MAPK signaling pathway.

### 2. The cilia phenotype in FSTL1 KD cells is reversed by inhibiting ERK phosphorylation

To further analyze the correlation between pERK and cilia length we used MAP Kinase MEK1 inhibitor U0126 to modulate the levels of pERK. In hTERT-RPE1 cells, ciliogenesis is induced by incubating confluent cells with serum free media for 24 hours. Inhibition of ERK phosphorylation was achieved by adding U0126 during this incubation. In cells transfected with control siRNA, treatment with U0126 resulted in downregulated ERK phosphorylation (Fig. 2A) and longer cilia than in control cells (2.97 μm compared to 2.68 μm in control cells) (Fig 2B). Consistent with our previous results, transfection with siRNA FSTL1 increased pERK a 2.5-fold (Fig. 2A) while the addition of U0126 to this condition, was able to fully reverse the effect of FSTL1 KD on both ERK phosphorylation and cilia length (from 1.73 μm to 2.47μm).

**Figure 2.**
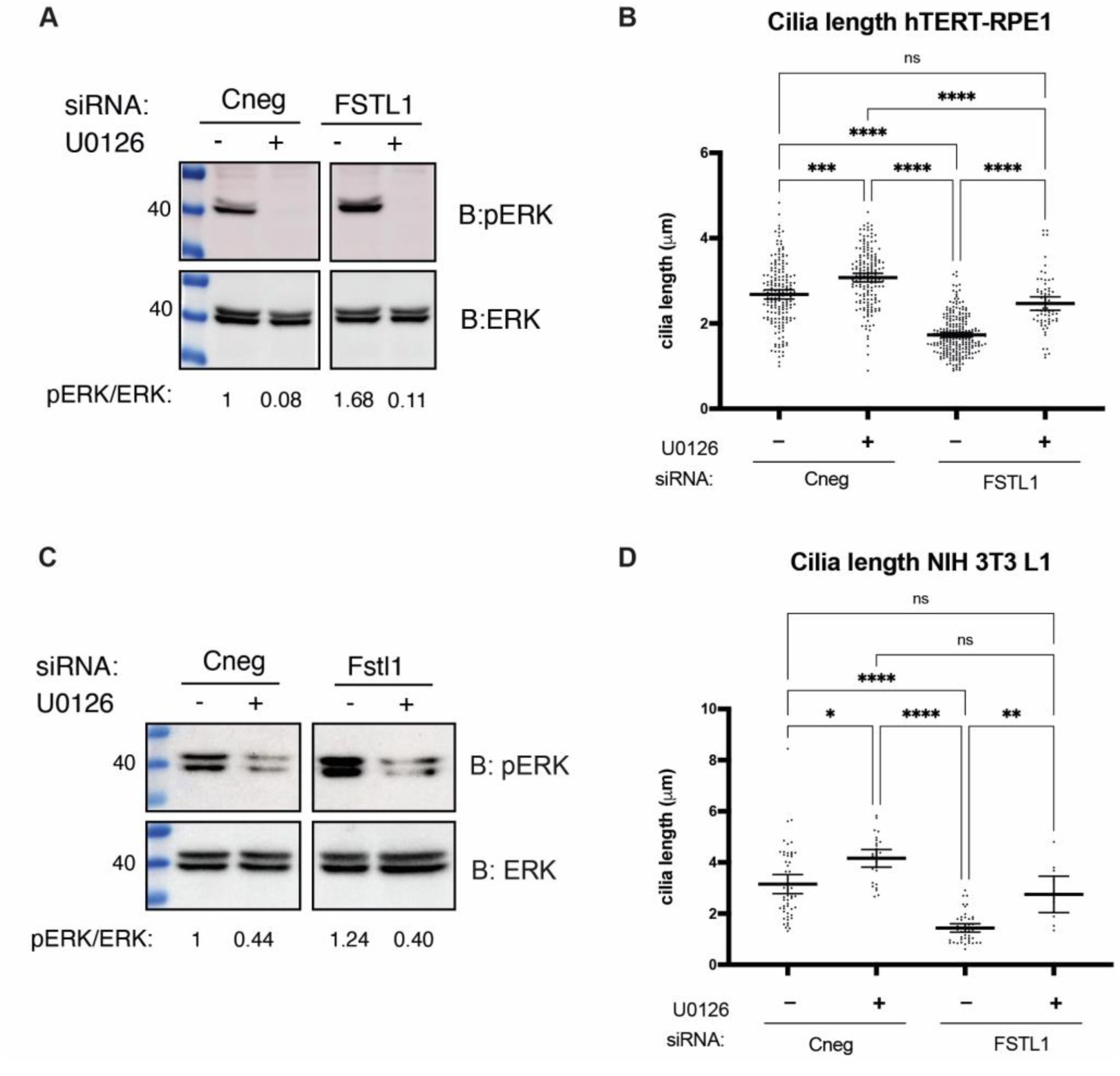
Inhibition of ERK phosphorylation reverses the effect of FSTL1 KD on cilia length. Cells were transfected with siRNA Cneg or siRNA FSTL1 for 48 hours. In the last 24 hours cell culture media were changed to FBS-medium supplemented with 40 μM U0126. hTERT-RPE1 (A) and 3T3-L1 (C) whole cells lysates were analyzed by Western blot anti-pERK and anti-ERK. Densitometric readings were used to calculate the ratio pERK/ERK. hTERT-RPE1 (B) and 3T3-L1 (D) cells were prepared for immunofluorescence anti-acetylated tubulin and anti-gamma tubulin and primary cilia were measured in each condition. The cilia length from each condition were measured and compared using a Kruskall-Wallis test. Dot plots with a line at the mean are shown and error bars represent 95% confidence intervals. Significant differences are indicated in comparison to control cells (siRNA Cneg; untreated) except when indicated with a bracket and are signaled with ns: not significant; *: P=0.01-0.05; **: P=0.001-0.01; ***: P=0.0001-0.001; ****: P<0.0001. Data shown are representative of at least 2 experiments each.

To expand on these results and to explore a link with a physiological process, we also tested the effect of pERK modulation in the preadipocyte 3T3-L1 cell line. Consistent with data shown in hTERT-RPE1 cells, Fstl1 knockdown increased pERK (Fig. 2C) and resulted in shortened cilia (1.42 μm compared to 3.15 μm in control cells) (Fig 2D). Again, treatment of control cells with U0126 produced inhibition of pERK and longer cilia (4.16 μm), and ERK inhibition rescued the cilia phenotype (2.75 μm) in FSTL1-KD cells. Thus, these data support the notion that ERK activation is a mediator in the role of FSTL1 in cilia length regulation.

Furthermore, we next explored the possibility that FSTL1 signaling involves the inhibition of ERK phosphorylation by acting as an antagonist of MAPK activating pathways.

Since FSTL1 is a known antagonist of BMP signaling, we analyzed whether the BMP canonical SMAD-mediated signaling pathways is involved in FSTL1 signaling in hTERT-RPE1 cells. However, pSMAD signaling is inhibited by the incubation with U0126 which suggests the existence of a crosstalk between ERK and SMAD pathways and by the incubation with recFstl1 suggesting an antagonic effect of FSTL1 on BMP/SMAD signaling. Knocking down FSTL1 did not modify the pattern of SMAD phosphorylation. Therefore, our data suggest that the absence of FSTL1 due to the siRNA FSTL1 does not affect SMAD phosphorylation meaning that SMAD pathway is not involved in the signaling by extracellular FSTL1 (Fig. S1).

### 3. ERK phosphorylation is required for cilia reabsorption during 3T3-L1 differentiation

Primary cilia dynamics is coordinated with preadipocyte differentiation whereby cilia initially extend during the first 2-3 days post-induction and then are reabsorbed and virtually absent of extremely short in fully differentiated adipocytes [25, 33]. In addition, the presence of Fstl1 in the media during differentiation blocks cilia reabsorption keeping cilia longer and also preventing a full differentiation [13].

Our data so far indicated that FSTL1 regulates cilia, both in RPE and 3T3-L1 cells, at least in part through modulating ERK phosphorylation. As mentioned, FSTL1 levels are high at the onset of differentiation and they are significantly reduced as adipogenesis proceeds [12, 13]. Also pERK levels peak around days 4-8 of differentiation and inhibition of pERK with U0126 promotes adipogenesis in 3T3-L1 cells, suggesting that Fstl1 role in adipogenesis is mediated by modulation of pERK [14]. Therefore, we focused on testing whether pERK inhibition would inhibit cilia reabsorption and subsequently adipogenesis, mimicking the effect of adding recombinant FSTL1.

We first incubated differentiating preadipocytes with 40 μM U0126 starting at day 0 with the differentiation cocktail and keeping it present in the media changes every 2 days. In the presence of U0126, differentiation of 3T3-L1 cells was inhibited compared to control cells incubated with the vehicle DMSO (Fig. S2). Also, in treated cells, we noted in the immunofluorescence staining of acetylated tubulin that the microtubule cytoskeleton was affected, possibly reflecting cellular stress. We therefore turned to shorter incubations to minimize pleiotropic effects of U0126 due to toxicity.

Next, we incubated differentiating cells for a 48-hour pulse between day 3 and day 5 post-induction and observed lipid droplets accumulation at different time points to assess the progression of differentiation at day 3, day 4 and day 10. At day 4, U0126-treated cells show a marked reduction in droplets accumulation compared to control cells which was however reversed at day 10 when both conditions did not differ in the observed lipid droplets accumulation (Fig 3A). We then added U0126 to cells undergoing differentiation at day 3 and measured ERK phosphorylation and cilia length before treatment (day 3) and after 24 hours with U0126 (day 4). We found that in control cells, ERK phosphorylation was slightly increased or maintained between day 3 and day 4 (Fig 4B) and we observed evidence of cilia reabsorption: cilia length was reduced at day 4 (1.9 μm at day 3 compared to 1.6 μm at day 4) (Fig 4C). In contrast, in cells treated with U0126, pERK was downregulated at day 4 and cilia reabsorption was impaired (averaging 2.1 - 2.0 μm at day 3 and day 4 respectively). Again, we observed diminished lipid accumulation at day 4 post-induction in U0126-treated cells compared to control cells (Fig 3D). In U0126-treated cells which had low levels of pERK and longer cilia we also observed a downregulation at day 4 of the expression of Pparγ (Fig 4E). Hence, although it does not inhibit differentiation at final time, short term incubation of U0126 after day 3 inhibits ERK phosphorylation and primary cilia reabsorption which may cause a delay in differentiation at day 4. These results suggest that primary cilia reabsorption mediated by Fstl1 downregulation during adipogenesis would contribute to promoting the progression of differentiation keeping high levels of pERK.

**Figure 3.**
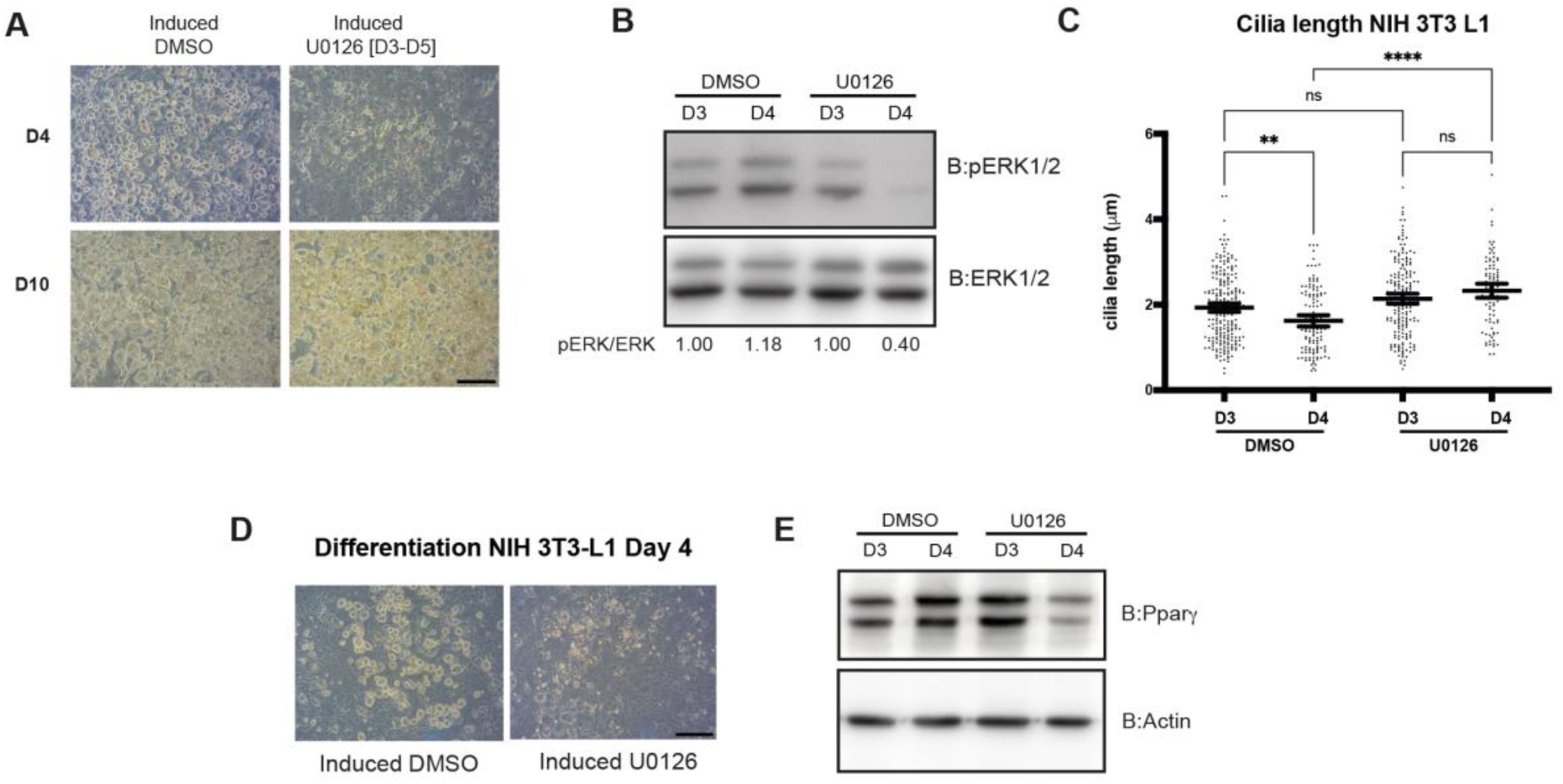
Short-term incubation with U0126 delays 3T3-L1 differentiation. (A) micrographs taken at D4 and D10 of cells induced to differentiate and incubated with a 48-hour pulse of U0126 between D3 and D5. Scale bar represents 100 μm. (B) Western blot showing ERK phosphorylation at D3 and D4 of differentiation after addition of MEK1 kinase inhibitor U0126 or control vehicle DMSO at D3. (C) cilia length was measured in each condition. (D) micrographs comparing lipid droplets accumulation at D4 in cells uninduced and induced cells incubated after D3 in the presence of U0126. Scale bar represents 100 μm. (E) Western blot showing Pparγ2 (higher MW band) and Ppraγ2 (lower MW band) expression at day 3 and day 4 after induction in the absence or presence of U0126. Addition of U0126 for 24 hours downregulates Pparγ expression. Data shown is representative of 3 experiments for A, D and E and of 2 experiments for B and C.

**Figure 4.**
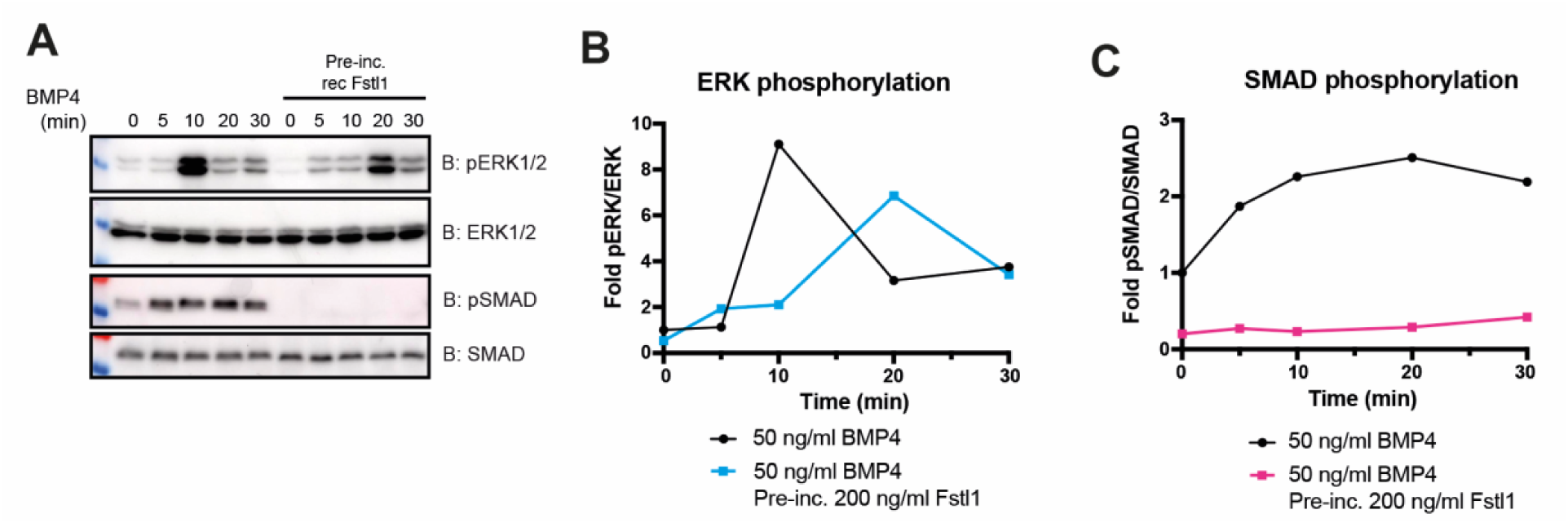
FSTL1 inhibits BMP4 non-canonical signaling. Western blot showing the time course of pERK and pSMAD after stimulation with BMP4. Confluent hTERT-RPE1 cells were serum-starved for 24 hours, incubated with 200 ng/ml Fstl1 for 1 hour and then stimulated with 10 ng/ml. Cells were collected at 0, 5, 10, 20 and 30 minutes after stimulation and analyzed by western blot anti-pERK, anti-pSMAD, anti-ERK and anti-SMAD. Densitometric readings were used to calculate ratios of pERK/ERK and pSMAD/SMAD. (B) Graph representing the time-course fold pERK/ERK comparing Fstl1-preincubated cells (blue line) with control cells only stimulated with BMP4. (C) Graph representing the time-course fold pSMAD/SMAD comparing Fstl1-preincubated cells (magenta line) with control cells only stimulated with BMP4. Preincubation with Fstl1 inhibits SMAD mediated signaling and delays peak phosphorylation of ERK.

### 4. FSTL1 modulates ERK-mediated, non-canonical BMP4 signaling in hTert-RPE1 cells

FSTL1 has been shown to interact in vitro with members of the TGFβ and BMP family [19] and reported to antagonize specifically BMP4 canonical signaling during lung development [3] and cervical carcinogenesis [7]. In addition, BMP4 and other BMP proteins can induce a non-canonical signaling pathway that involves MAP Kinases [34, 35]. Therefore, we asked whether FSTL1 could affect ERK phosphorylation by interfering with BMP4 non-canonical signaling.

Consistent with previously reported data [36] the incubation of serum-starved hTERT-RPE1 cells with recombinant BMP4 activates the phosphorylation of ERK and SMAD with specific time-course dynamics (Fig. 4A)

The stimulus using 10 ng/ml of recombinant BMP4 caused a fast increase of pSMAD after 5 minutes remaining high until at least 30 minutes of incubation while pERK1/2 showed a peak at 10 minutes and was down regulated after 20 minutes (Fig. 4A, lanes 1-5). We then, pre-incubated the cells with 200 ng/ml of recFstl1 for 1 hour before BMP4 stimulation. First, the pre-incubation with FSTL1 was able to fully abrogate SMAD phosphorylation thus indicating inhibition of canonical BMP4 signaling (Fig. 4A. lanes 6-10). In the case of ERK, basal phosphorylation levels (0 min) were also reduced due to FSTL1is case and the peak of ERK phosphorylation was of lower intensity and delayed (detected at the 20 min time point; Fig. 4A, lanes 6-10). These data suggest that FSTL1 interferes in a competitive manner with BMP4 not only in the context of canonical-SMAD signaling but also non canonical BMP4 signaling involving ERK phosphorylation. Taken together, our data suggest that the mechanism by which FSTL1 regulates ciliogenesis involves ERK phosphorylation, possibly downstream of a receptor of the BMP family.

## Discussion

The secreted glycoprotein FSTL1 is emerging clearly as an important factor in multiple processes involved in development and pathogenesis. While FSTL1 involvement in organogenesis, cancer, cardiovascular disease, inflammation, and obesity is clear, the detailed molecular mechanisms for FSTL1 signaling remain to be elucidated. A detailed revision of FSTL1 molecular mechanisms of action suggests a tissue-dependent action for FSTL1 [1]. One source for this variation might be related to the receptor that would be targeted by FSTL1 in each situation. For example, DIP2A and BMP receptors were reported to act upstream of FSTL1 in cardiovascular disease, activating AKT [22] and SMAD [37] signaling pathways respectively. Other intracellular mediators such as ERK, AMPK have been identified as responding to FSTL1 in this cell type, although their receptors are unknown [1]. During lung organogenesis, Fstl11 antagonizes BMP4 canonical signaling mediated by SMAD proteins leading to lethal lung malformation [3]. In cancer biology, several signaling pathways have been shown to respond to FSTL1 both in cell growth and metastasis and FSTL1 has been reported to play both pro- and anti-tumorigenic roles. Overall, multiple mechanisms seem to mediate cancer development although only a few receptors have been proposed in these pathways (e.g. Fas-Ligand receptor and Connexin 43). Similarly, the knowledge on inflammation is currently fragmented and the mechanisms involved in FSTL1 signaling seem to differ in a tissue- and pathology-specific manner. Importantly, the glycosylation state of the N-glycosylation sites of FSTL1 is essential for biological activity and different effects have been reported depending on this posttranslational modification [4, 14, 38].

In addition to the above-mentioned functional roles of FSTL1 [1], we have previously uncovered a novel role for FSTL1 as a ciliogenesis regulator whereby knocking down FSTL1 in hTERT-RPE1 cells, decreases primary cilia length. In turn, adding back Fstl1 in the cell culture media rescues the ciliary length [13]. This effect is recapitulated in murine preadipocytes 3T3-L1 and we found that modulation of cilia length by Fstl1 in the extracellular medium regulates adipogenesis in culture. Moreover, ERK modulates ciliogenesis [27, 28] and Erk phosphorylation is regulated by Fstl1 during adipocyte differentiation [14].

Thus, we have developed a working model by which FSTL1 regulates cilia-mediated signaling pathways that in turn regulate biological processes such as adipogenesis. In this work we aimed to determine which signaling pathway mediates the role of FSTL1 in ciliogenesis and adipogenesis.

Here, we provide evidence for ERK as mediator of ciliogenesis downstream of FSTL1 and we provide data that suggests a molecular mechanism by antagonizing BMP4 signaling. In addition, we add new evidence linking the ciliogenesis regulation by Fstl1 to Erk phosphorylation during adipogenesis.

Following up on our previous findings of the role of FSTL1 in ciliogenesis, we turned to investigate known binding partners of FSTL1. These include several members of the TGFβ family such as BMP4, TGFβRII and BMPRI, as well as DIP2A [19]. We set out to analyze the role of these primary candidates in FSTL1 signaling. By depleting FSTL1 in cell culture, we found that FSTL1 antagonizes ERK phosphorylation (Fig1A). In turn, we found that ERK phosphorylation dynamics correlates with primary cilium length (Fig.1 B-E). This suggests that the regulation of ciliogenesis by FSTL1 is mediated by the MAP Kinase pathway via ERK phosphorylation. This finding is consistent with the previous reports of MAP Kinases linked to ciliogenesis and cilia length regulation. Cisplatin treated cells kidney tubular epithelial cells have shorter cilia and higher pERK while inhibition of pERK by U0126 during cisplatin treatment preserves cilia [28]. ERK is primarily regulated by MEK1 kinase and by DUSP6 phosphatase. Interestingly, the activation of ERK by a DUSP6 inhibitor reduces cilia length in Chlamydomonas reinhardtii and in hTERT-RPE1 cells [27]. We also tested any effect of FSTL1 depletion on SMAD1/5/8 activation as a mediator of the canonical BMP signaling pathway. At this point we cannot rule out a role for DIP2A since we were not able to test it in ciliogenesis, but future efforts should be pointed to clarify the role of DIP2A in ciliogenesis and possibly in adipogenesis. Also, in our hands, we did not observe any variation on phospho-SMAD1/5/8 content upon FSTL1 depletion and inhibition of ciliogenesis, suggesting that these pathways are not involved in FSTL1 signaling in hTERT-RPE1 cells. However these findings do not exclude a TGFβ/BMP receptor mechanism since a growing body of evidence shows that in addition to the canonical SMAD-mediated signaling pathway, BMPs are able to signal using a non-canonical pathway mediated by MAPK, including ERK [39]. Indeed, although the evidence available for this effect come from experiments that do not identify the initiation mechanisms, BMPs are able to activate another intracellular mediator including the MAP Kinase ERK, PI3K, PKA, PKC and PKD [40].

The primary cilium is a sensory organelle that is present in virtually every cell in a multicellular organism, and we had shown that depletion of FSTL1 leads to shorter cilia. We tested the involvement of pERK in the cilia phenotype caused by FSTL1 knock down using U0126 to inhibit pERK in the presence or absence of FSTL1 both in human and murine cells. We find that inhibition of ERK phosphorylation rescues the cilium phenotype caused by FSTL1 knock down (Fig 2). Thus, to the previously known roles of ERK in cilia length regulation and of FSTL1 in ERK phosphorylation, we contribute that FSTL1 controls cilia length at least partially through the modulation of ERK phosphorylation.

Defects in ciliary structure or function have been described to be associated to a set of genetic mutations and cause a group of pathologies called ciliopathies [23]. Among ciliopathies, Alström Syndrome (ALMS) and Bardet Biedl Syndrome (BBS) are diseases characterized by multiple symptoms that include obesity in patients as well as in animal transgenic models [23, 41–45].

Some of the signaling pathways during adipogenesis include the anti-adipogenic Wnt and Hedgehog pathways -which are known to be coordinated by primary cilia- and pro-adipogenic signals such as IGF1, FGF, Activin, TGFβ, BMP [26, 46]. Consistent with a previous report that proposed FSTL1 as a biomarker for adipogenesis [12], we described a role for Fstl1 modulation in 3T3-L1 preadipocyte differentiation in culture. We found that Fstl1 is required at the onset of adipogenesis and then it is required to be downregulated to allow for differentiation to proceed. Artificially keeping high levels of Fstl1 prevented full differentiation of 3T3-L1 [13]. In parallel, we and others observed that primary cilia are present at the onset of differentiation, elongate at day 2 after inducing differentiation and are reabsorbed as the process is completed [13, 25] Since inhibition of ERK phosphorylation in 3T3-L1 preadipocytes correlates with longer cilia we asked whether pERK regulation of cilia might be responsible for Fstl1 modulation of adipogenesis. Therefore, we next tested whether regulation of pERK by Fstl1 affects primary cilia during 3T3-L1 adipocyte differentiation.

We found that the inhibition of ERK phosphorylation during differentiation keeps the primary cilia longer and mimics the effect of keeping high levels of Fstl1 during adipogenesis (Fig. 3). We incubated preadipocytes already undergoing the first stages of differentiation (day 3) with a pulse-like single dose of U0126 which was able to inhibit ERK phosphorylation (Fig. 3B). We found that inhibition of pERK in early stages of differentiation interfered with the cilia dynamics preventing their reabsorption at day 4 (Fig. 3C). Pparγ expression is a transcription factor that is induced during adipogenesis (Fig. 3E) and plays a central role in controlling the expression of adipocyte specific genes [47]. We found that the pulse inhibition of pERK led to the downregulation of Pparγ (Fig.3E) which correlated to an apparent delay in lipid droplets accumulation (Fig. 3D), suggesting a partial reversion or delay in differentiation. These data are reminiscent to the effect of preventing Fstl1 downregulation during 3T3-L1 adipogenesis and therefore suggest that Fstl1 role in differentiation is mediated by pERK. These findings reinforce the interest on Fstl1 as a key player in adipogenesis and adipose tissue homeostasis. Consistent with our findings in cell culture [13], in vivo experiments confirmed that the differentiation to adipocytes of Fstl1-/- cells (MEFs of SVCs) derived from mice embryos was impaired [14].

Moreover, FSTL1 has been proposed as a potential biomarker in inflammatory diseases in mice [48] and specifically for metabolic health in obesity [16]. Future work will be focused on studying the role of Fstl1 in adipose tissue.

Information about mediators of FSTL1 signaling is provided by in vitro experiments of FSTL1 binding to members of the TGFβ family and other ligands. These studies show that FSTL1 has the potential to bind to DIP2A, BMP2 and BMP4 and other members of the superfamily such as BMPR1A, TGFβ and activin [3, 19]. FSTL1 has been reported as an antagonist of BMP signaling in several experimental models. During zebrafish development Fstl1a and fstl1b antagonize SMAD-mediated BMP4 activity as depletion of fstl1 increases the levels of pSMAD1/5/8 and interferes with the correct maturation of the notochord [49]. In the context of mouse lung development, Fstl1 inhibits BMP4/SMAD1/5/8 signaling [3] while FSTL1 contributes to the pathogenesis of cervical cancer by negatively regulating BMP4/SMAD1/5/8 signaling [7]. In pulmonary artery smooth muscle cells the antagonistic effect of FSTL1 on BMP4 is mediated by downregulating pERK [32] as an example of non-canonical signaling pathways activated by TGFβ/BMPs [35].

We investigated how FTSL1 would impact on BMP4 signaling in our ciliogenesis model. First, we found that in hTERT-RPE1 cells BMP4 activates the pERK as well as pSMAD1/5/8. Consistent with studies on BMP9 and BMP2 in the osteoblast precursor cells line MC3T3-E1 [36], this activation is transient and time dependent (Fig. 4A). When the BMP4 treatment is performed in cells pre-incubated with an excess of recombinant fstl1, we find that Fstl1 non only attenuates SMAD activation as previously described but also interferes in ERK activation by BMP4. These results suggests that in hTERT-RPE1 cells, FSTL1 regulates ciliogenesis by antagonizing BMP4 non canonical signaling mediated by ERK. We did observe that FSTL1 inhibits BMP4/SMAD-mediated canonical signaling although thus far we do not have evidence that SMAD controls ciliogenesis (Fig. S1)

These data might be relevant to the regulation adipocyte differentiation by FSTL1/ERK/cilia that we propose here (Fig. 3) since BMP proteins are involved in adipogenesis and specifically BMP4 preadipocyte commitment [47, 50, 51]. Further studies will be needed to elucidate the role of FSTL1-regulated ciliogenesis in adipogenesis and adipose tissue biology. In that sense, BMP4 as well as BMP2 were shown to be required for adipocyte lineage commitment in a model of pluripotent cells C3H10T1/2 [52]. During adipogenesis and also terminal differentiation to brown adipose tissue BMP4 binds to its heterodimeric receptor formed by BMPR1A or BMP1RB (Type I) and BMPr2 (Type II) [50]. In a systemic level, high serum concentrations of BMP4 are associated with obesity and Type-2 Diabetes [53] and recent in vivo studies have shown that BMPr2 is required for fat accumulation, possibly by promoting adipocyte survival [54].

Future work will be aimed at identifying the receptor acting upstream of FSTL1 in ciliogenesis and adipogenesis regulation. Based on our data this receptor is most likely a BMP4-binding protein. Since BMPs bind to dimeric BMP receptors, candidates include ALK and BMPr2 as well as other members of the TGFβ family. It also remains to be discovered which effector molecules mediate ERK phosphorylation and cilia length regulation. Some evidence point to actin cytoskeleton regulator and future studies should also include the exploration of this possible pathway [55, 56]. Other possible candidates as effectors of FSTL1 in ciliogenesis are MEK1 and DUSP6, which control the phosphorylation balance of ERK.

Therefore, we propose a mechanism based on BMP4 non-canonical signaling for the role of FSTL1 in ciliogenesis and adipogenesis in which FSTL1 antagonizes ERK activation, thereby regulating ciliogenesis and possibly cilia-supported anti-adipogenic signaling pathways.

## Materials and methods

### Reagents, antibodies

Cell culture media DMEM glutamax and DMEM F12, cell culture grade reagents PBS, antibiotics, glutamine, Hepes were from Gibco. Bovine serum albumin, TRIS buffer and salts for PBS, Tween-20, Sigmafast protease inhibitor cocktail tablets, Rosiglitazone and Insulin from Sigma; Triton X-100, Dexamethasone and 3-Isobutyl-1-Methylxanthine (IBMX) from AppliChem. Recombinant Fstl1, recombinant BMP4 from R&D, U0126 was from Promega (V1121) or Abcam (AB120241). Stealth RNA for FSTL1, control scrambled si RNA Low GC and Lipofectamine RNAiMAX were from Invitrogen (Thermo). siRNA smart pool of 4 oligos for BMPr2 was from Dharmacon. Antibodies mouse anti-Acetylated tubulin, rabbit anti gamma-tubulin, anti-alpha tubulin were form Sigma; anti-human FSTL1 (AF1694), anti-mouse Fstl1 (AF1738) were from R&D; rabbit mAb anti-phosphoERK1/2 (clone D13.14.4E), rabbit mAb anti-p42/44 MAPK (Erk1/2) (clone 137F5), rabbit mAb anti-phosphoSMAD1/5/9 (clone D5B10), anti-rabbit mAb SMAD1 (clone D59D7), anti Pparγ (clone 81B8) were from Cell Signaling; conjugated antibodies anti-mouse IgG-AlexaFluor488 (A21202), anti-rabbit IgG-AlexaFluor633 (A21070), DAPI was from Thermo (D21490).

Horseradish peroxidase (HRP)-conjugated anti-rabbit from Sigma (A0545) and anti-goat from Thermo (A16005).

### Immunofluorescence and imaging

A coverslip was added to the plates where cells were grown, differentiated, or treated with inhibitors. Coverslips with adhered cells, were fixed in ice-cold methanol 10 min at -20°C, permeabilized with 0,1% Triton X-100 in PBS and blocked with 0,5% SBF in PBS 1 hour at room temperature. Coverslips were then incubated for 1 hour at room temperature with the corresponding primary antibody (anti-acetylated tubulin and anti-gamma tubulin, each diluted 1/1000 in PBS), then washed three times briefly with PBS and incubated for 1 hour at room temperature with conjugated antibodies and DNA-staining dye (anti-mouse AF 488, anti-rabbit AF 633 each 1/1000 and DAPI (6 μg/ml). After 3 washes, coverslips were mounted in Prolong antifade. Images of cilia were obtained using a confocal microscope Zeiss 880 using a 63X objective and 1.4 refraction index immersion oil. Cells stained with Oil Red O were imaged using an inverted microscope Zeiss Primovert for transmitted light with 20X or 40X objectives and coupled to a color digital camera Axiocam ERc 5s.

### Cilia measurements and statistical analyses

Confocal images of cilia were processed and analyzed, and individual cilia lengths were measured using the freehand ROI selection tool in FIJI. All data was analyzed using GraphPad Prism 10. Al least 50 (and up to 150) cilia were measured for each condition. Western blots were analyzed using the densitometry tools form FIJI and data were also analyzed using Prism 10. Following outliers’ identification and normality tests, all data were analyzed using T-test or Kruskall-Wallis test with Dunn’s test for multiple comparisons. In all cases, P-values <0.05 were used to conclude on significant differences.line

### Western blotting

Cells were collected, washed once with PBS and then lysed using 50 mM Tris-HCl pH7.5, 50 mM NaCl, 1 % Nonidet P40. Cells were incubated 45 min at 4°C, centrifuged at 12000 g and supernatants were recovered as the whole cell lysate (WCL). Protein concentration of WCL was measured using Bincinchonic acid assay (Pierce) and 30-50 μg of protein was loaded per well in 10% SDS-PAGE gels. After electrophoresis, the proteins were transferred to a PVDF membrane (-), blocked with 5%BSA in PBS-0.1% Tween-20 and incubated overnight at 4°C with primary antibodies (anti-pERK, anti pSMAD1/5/9, anti-ERK, anti-SMAD1), washed and then incubated with Horseradish peroxidase-conjugated antibodies. For detection, membranes were incubated for 1-5 min with ECL reagent and the Western blot images were captured using the Imagequant800 system (Amersham). Densitometric readings were obtained using FIJI and used to calculate different ratio values.

### Cell culture and adipocyte differentiation

hTERT-RPE1 cells were cultured in DMEM-F12 media supplemented with 10% FBS, Hepes 0,01 mg/ml Hygromycin (Sigma) and 100 U/ml Penicillin/100 μg/ml Streptomycin (PS). Cilia growth was induced by culturing cells to a high confluency followed by 24 hours serum starvation. -3T3-L1 cells were cultured in DMEM-glutamax high glucose high pyruvate, supplemented with 10% FBS Hepes and PS. For adipocyte differentiation -3T3-L1 cells were seeded at 0.8-1x10^6^ cells/well, left 2 days after confluence, treated for 48hs with cocktail differentiation (DMEM with 10% FBS, PS, 1 μg/ml bovine insulin, 0.25 mM dexamethasone, 0.5 mM IBMX and 2.5 mM Rosiglitazone) and then the medium was changed every two days with DMEM with 10% FBS, PS, 1 μg/ml insulin during 8 additional days.

### Gene silencing

Gene silencing of human and murine FSTL1 was achieved as described [13]. Briefly, cells were transfected with stealth siRNA duplex ARN oligos (human: GGAACAGAAUGAAACUGCCAUCAAU; mouse:

CCUAGACAAGUACUUUAAGAGCUUU) using lipofectamine RNAiMAX.

### U0126 treatments and stimulations

To inhibit ERK1/2 phosphorylation, cells were incubated in fresh medium supplemented with 40 μM U0126 for 20-24 hours and then harvested for Western blotting or immunofluorescence. Recombinant Fstl1 was added to cells at 200 ng/ml with fresh medium and incubated overnight for rescue experiments or for 1 hour for competition experiments. Recombinant BMP4 was added to 24hs serum-starved cells at 10 ng/ml for different time points between 5 minutes and 30 minutes.

## Supplementary data

**Figure S1.**
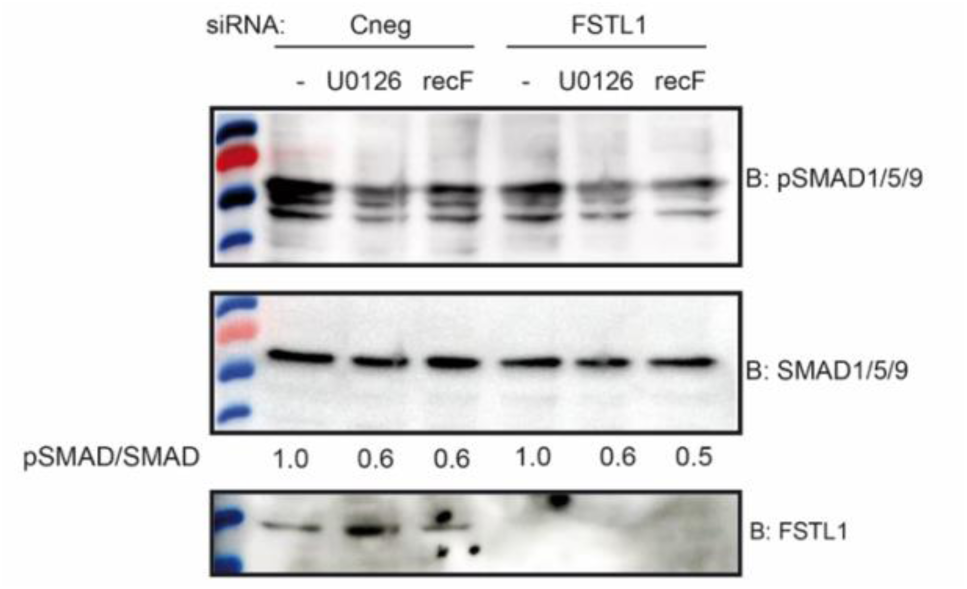
FSTL1 does not signal using SMAD pathway. hTERT-RPE1 cells were transfected with siRNA Cneg or siRNA FSTL1 for 48 hours. In the last 24 hours cell culture media were changed and FBS-medium was added with or without supplementation with 200 ng/ml recFstl1. Cell lysates were analyzed by Western blot anti-pSMAD, anti-SMAD and anti-FSTL1. Densitometry readings were used to calculate pSMAD/SMAD ratios with each condition normalized to the siRNA Cneg transfected and untreated cells. The absence of FSTL1 does not modify SMAD phosphorylation in any of the treatments.

**Figure S2.**
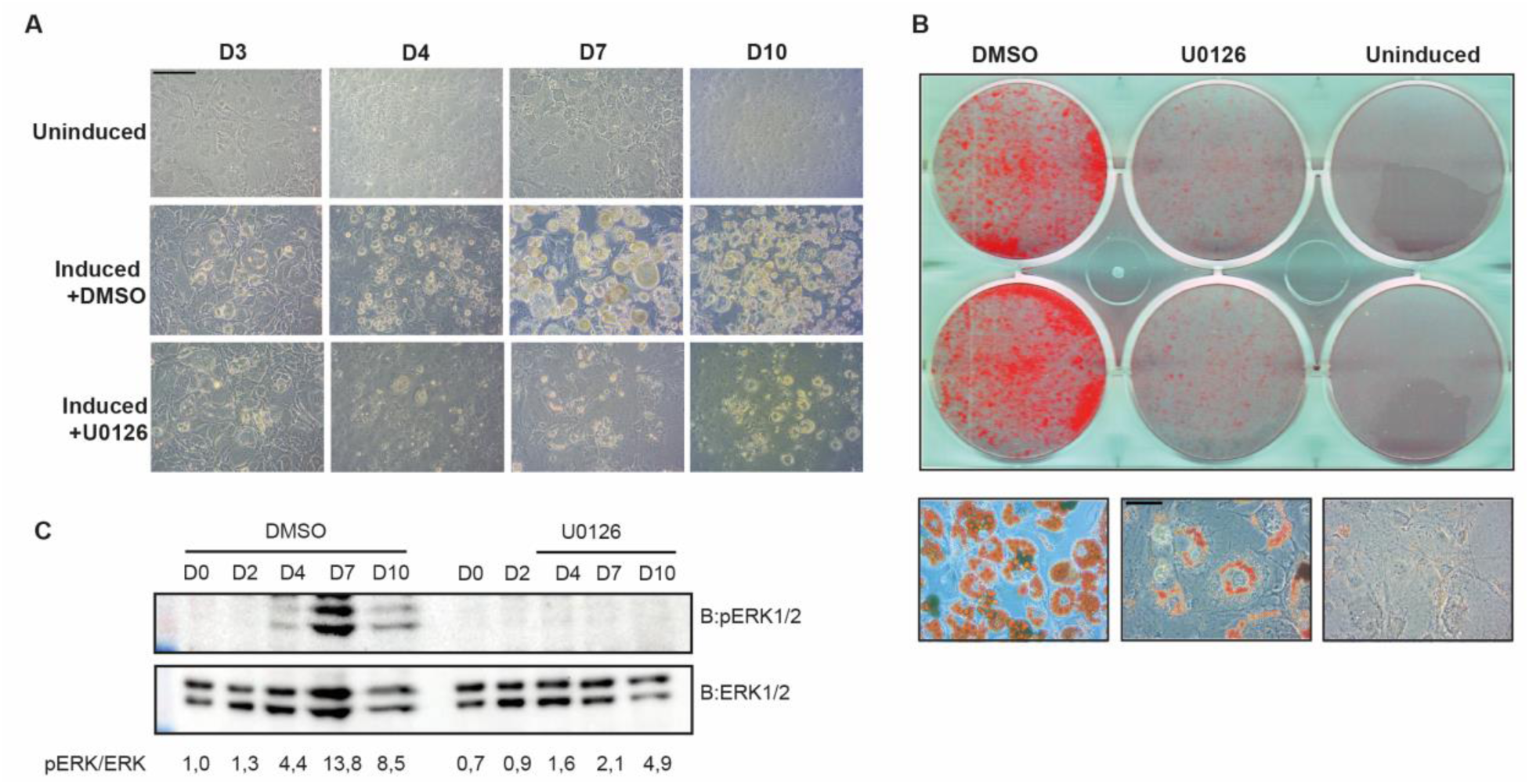
U0126 inhibits 3T3-L1 differentiation. 3T3-L1 cells were induced to differentiate in the presence of 40 mM U0126 staring at Day 0 and changing every 2 days. (A) Micrographs of live cells uninduced control or induced to differentiate in the presence of U0126 or control vehicle DMSO. Images were taken at different time points during differentiation day 3, day 4, day 7 and day 10. Scale bar represents 50 mm (B) Oil Red O staining comparing lipid accumulation at day 10 in uninduced control or induced to differentiate in the presence of U0126 or control vehicle DMSO (top panel) and micrographs of Oil Red O staining showing a closer image (40X) of stained lipid droplets (bottom panel). Scale bar represents 10 mm (C) Western blot showing time course phosphorylation of ERK during differentiation in the absence (left) or presence of U0126 (right).

## Acknowledgments

We thank Paola Lepanto, Magdalena Cárdenas-Rodríguez, Florencia Irigoín, Ileana Sosa for discussions along different stages of this work. We thank Dr. Luisa Berná for critically reading and help editing the manuscript. We acknowledge the Advanced Bioimaging Unit at the Institut Pasteur de Montevideo for their support and assistance.

## Authors contribution

LS. Design and performing the experiments and data analysis, manuscript revision; LG. Performed experiments and analyzed data, manuscript revision; CE. Discussion and data analysis, manuscript revision and edition; JLB. Project coordination, experiment design, discussion and data analysis manuscript revision and edition; VPE. Funding acquisition, project coordination, experiment design, discussion and data analysis, Manuscript drafting and writing.

